# Tn-seq of the *Candida glabrata* reference strain CBS138 reveals epigenetic plasticity, structural variation, and intrinsic mechanisms of resistance to micafungin

**DOI:** 10.1101/2024.05.02.592251

**Authors:** Timothy J. Nickels, Andrew P. Gale, Abigail A. Harrington, Winston Timp, Kyle W. Cunningham

## Abstract

*C. glabrata* is an opportunistic pathogen that can resist common antifungals and rapidly acquire multidrug resistance. A large amount of genetic variation exists between isolates, which complicates generalizations. Portable Tn-seq methods can efficiently provide genome-wide information on strain differences and genetic mechanisms. Using the *Hermes* transposon, the CBS138 reference strain and a commonly studied derivative termed 2001 were subjected to Tn-seq in control conditions and after exposure to varying doses of the clinical antifungal micafungin. The approach revealed large differences between these strains, including a 131 kb tandem duplication and a variety of fitness differences. Additionally, both strains exhibited up to 1000-fold increased transposon accessibility in subtelomeric regions relative to the BG2 strain, indicative of open subtelomeric chromatin in these isolates and large epigenetic variation within the species. Unexpectedly, the Pdr1 transcription factor conferred resistance to micafungin through targets other than *CDR1*. Other micafungin resistance pathways were also revealed including mannosyltransferase activity and biosynthesis of the lipid precursor sphingosine, the drugging of which by SDZ 90-215 or myriocin enhanced the potency of micafungin *in vitro*. These findings provide insights into complexity of the *C. glabrata* species as well as strategies for improving antifungal efficacy.

**Summary:** Candida glabrata is an emerging pathogen with large genetic diversity and genome plasticity. The type strain CBS138 and a laboratory derivative were mutagenized with the *Hermes* transposon and profiled using Tn-seq. Numerous genes that regulate innate and acquired resistance to an important clinical antifungal were uncovered, including a pleiotropic drug resistance gene (PDR1) and a duplication of part of one chromosome. Compounds that target PDR1 and other genes may augment the potency of existing antifungals.

## INTRODUCTION

Many species of single-cell fungi live commensally with their human hosts. Several of these species can penetrate epithelial barriers and cause disease, especially in immunocompromised individuals (Yang et al. 2014). Although *Candida albicans* is the most common causative agent of candidiasis, the prevalence of *Candida glabrata* infection has been rising compared to *C. albicans* and other causative species (Guinea 2014; Pfaller et al. 2019) due to both intrinsic and rapidly acquired resistance to the different classes of antifungal drugs (Alexander et al. 2013; Angoulvant et al. 2016; Pfaller et al. 2012; Singh-Babak et al. 2012). *C. glabrata* can cause many types of infection that vary in severity including vulvovaginal candidiasis, oral thrush, bloodstream (candidemia), and internal organs (invasive candidiasis). More severe infections are associated with lengthy hospitalizations and mortality rates as high as 60% (Gupta et al. 2015; Pappas et al. 2018; Pappas et al. 2003; Wisplinghoff et al. 2004). With a limited arsenal of antifungal drugs, understanding how these pathogens are developing resistance and identifying potential new targets for antifungal therapy is of the utmost importance.

The rapid evolution of pathogenicity and drug resistance in *C. glabrata* can be explained by its genetic plasticity. This species colonizes as an obligate haploid, and has only been observed undergoing asexual reproduction. Little evidence of a diploid form has been discovered, although rare mating events have been theorized due to the presence of distinct mating genes, mating types, and evidence of past recombination events during its evolution (Brisse et al. 2009; Carrete et al. 2018; Dodgson et al. 2005; Srikantha et al. 2003; Wong et al. 2003). As a result, the vast majority of genetic diversity between *C. glabrata* isolates comes from DNA replication errors, tolerance of DNA damage during replication (Shor and Perlin 2021), and chromosomal abnormalities or aneuploidies (Guo et al. 2020; Muller et al. 2009; Polakova et al. 2009; Zheng et al. 2022). These high rates of mutation and chromosomal rearrangements provide an explanation for *C. glabrata*’s increasingly successful colonization of its host in spite of advances in antifungal therapy. In agreement with this, many strains recovered from patients have mutations in a mismatch repair gene that leads to increased drug resistance when disrupted (Healey et al. 2016).

Genome sequences of clinical isolates reveal large amounts of variation at the nucleotide level as well as in chromosome structure (Biswas et al. 2018; Carrete et al. 2018; Guo et al. 2020; Helmstetter et al. 2022; Marcet-Houben et al. 2022; Xu et al. 2021) which define seven distinct clades that differ greatly from one another (Carrete et al. 2018). Most research has been performed on isolates from clades V and VII: CBS138 and BG2. The CBS138 strain was originally isolated from the feces of a healthy adult (Dujon et al. 2004). The BG2 strain was isolated from a patient with fluconazole-resistant vaginitis (Cormack and Falkow 1999). These two strains are 0.9% divergent from each other at the nucleotide level (Xu et al. 2021) and contain several chromosomal inversions and translocations with respect to each other (Marcet-Houben et al. 2022; Muller et al. 2009). They also are reported to have differences in metabolism and virulence (Usher et al. 2023) as well as a differential response to antifungal drugs which extends to other *C. glabrata* isolates (Ksiezopolska et al. 2021). In addition, laboratory derivatives of CBS138 have been found with altered karyotypes and length polymorphism on chromosome K, which was accompanied by increased resistance to the echinocandin-class antifungal drug caspofungin (Bader et al. 2012; Brunke et al. 2014). An undefined, yet stable, ∼130 kb duplication in chromosome K may cause these phenotypic changes (Bader et al. 2012; Brunke et al. 2014). Understanding the phenotypic, and underlying genetic, variation between isolates of *C. glabrata* and how it relates to drug resistance and virulence will lead to better antifungal therapies and patient outcomes.

Previous work has characterized essential genes, as well as genes important in resistance and susceptibility to the azole-class antifungal fluconazole in a BG2 strain derivative using a transposon mutagenesis technique (Gale et al. 2020). By generating new pools of transposon mutants in the CBS138 reference strain and the derived strain 2001, we define the boundaries of a tandem duplication on the left tip of chromosome K. This 131 kb tandem duplication is present in numerous commonly studied laboratory strains, including 2001HTL that is the parent strain of hundreds of individual gene knockout mutants (Schwarzmuller et al. 2014). Using Tn-seq, we document fitness differences between CBS138 and 2001 in addition to fitness differences relative to the BG2 strain. Remarkably, CBS138 and 2001 strains share enormously increased transposon accessibility within most subtelomeres, suggesting more open chromatin relative to BG2. Finally, we investigate intrinsic resistance to micafungin on a genome-wide scale and identify cellular processes and pathways that could potentially be drugged for enhancement of micafungin efficacy.

## RESULTS

### Transposon Insertion Sequencing reveals a tandem duplication in 2001u and most derivatives of CBS138

The *C. glabrata* species contains a large amount of genetic variation, with many isolates and clades exhibiting nearly 1% sequence variation from one another. To begin exploring the functional consequences of this variation, we mutagenized the genome of strain 2001u, a *ura3Δ* derivative of the highly studied 2001 strain (Kitada et al. 1995) that was thought to be identical to reference strain CBS138, using the same *NAT1*-marked *Hermes* transposon used previously to mutagenize strain BG2u (alias: BG14) (Gale et al. 2020). Two pools of insertion mutants in 2001u were generated independently and the insertion sites were PCR amplified, sequenced, mapped to the BG2 reference genome (Xu et al. 2021), and imaged using the IGV genome browser. The patterns and densities of insertions were similar for 2001u and BG14 throughout most of the genome. While the BG2 genome is 5.85% unmappable due to non-unique 75-mers, approximately 7.15% of the 2001 (CBS138) reference genome could not be mapped uniquely to the BG2 genome due to sequence variation between the two strains (Fig. 1A, unmappable). In spite of the lower mappability, two genomic regions exhibited noticeably more insertion sites in 2001u relative to BG2u: the subtelomeric regions of most chromosomes (see next section) and the left end of chromosome K (Fig. 1A).

**Figure 1.**
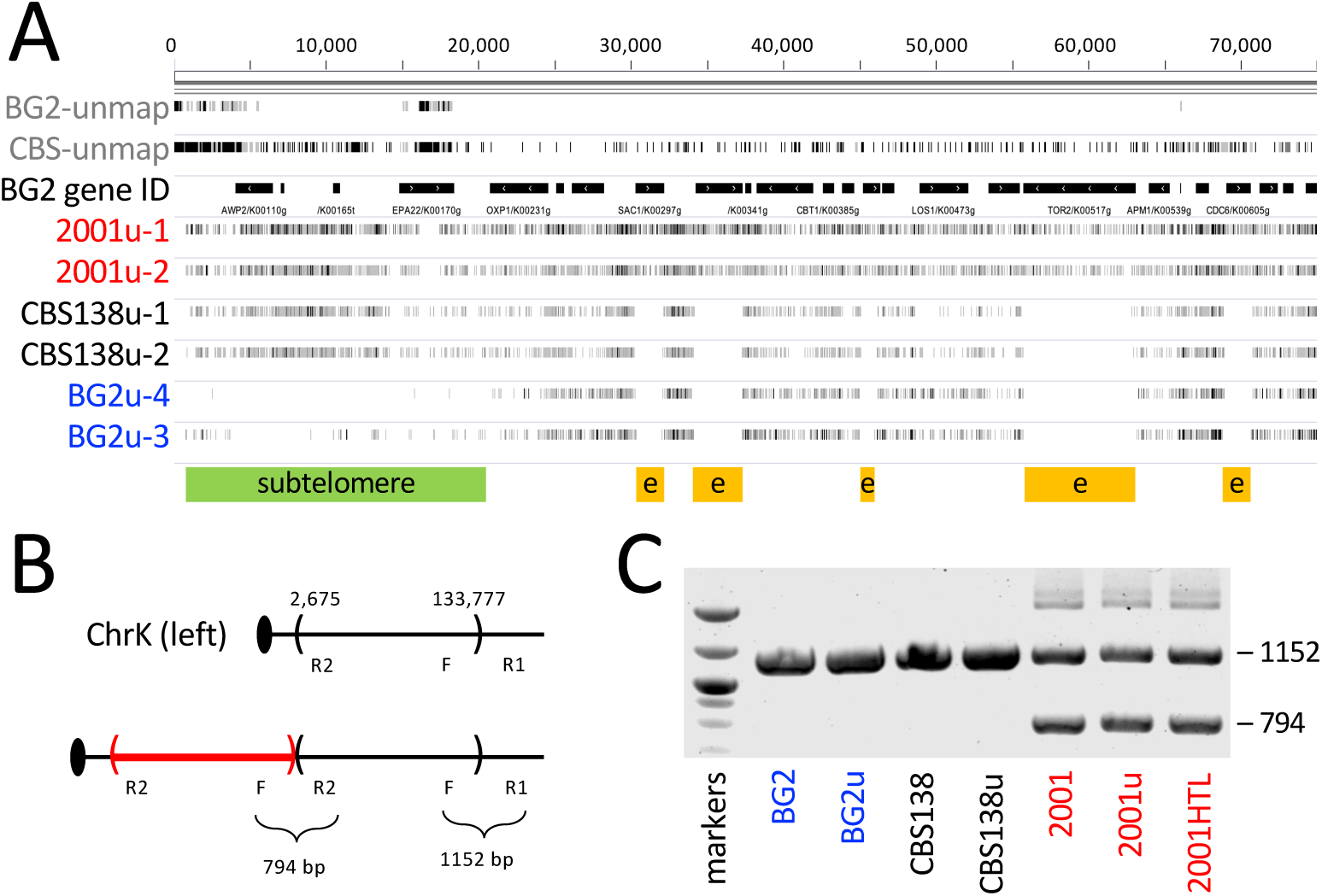
Detection of a large tandem duplication on the left tip of chromosome K in 2001-related strains. (A) A 75 kilobase segment of chromosome K of strain BG2 beginning with the telomere was visualized using the IGV genome browser including gene identifications (black bars), essential genes (orange bars), and the approximate position of the subtelomere (green bar). The positions of individual K-mers from the BG2 and CBS138 reference genomes that do not map uniquely to the BG2 reference genome (unmap) are indicated with gray or black tick marks depending on whether the K-mer was unmapped in one or both directions. Two independent pools of *Hermes* insertion mutants were generated in strains 2001u, CBS138u, and BG2u, sequenced, mapped to the BG2 reference genome, and visualize with gray to black tick marks at each insertion site. Darker tick marks indicate larger number of sequence reads mapping to each site. (B) Schematic of the left arm of chromosome K from CBS138u (top) and 2001u (bottom) illustrating a 131 kb tandem duplication (red) and the positions of forward (F) and reverse (R1, R2) primers that can diagnose its presence and absence. (C) Agarose gel analysis of genomic DNA prepared from BG2, CBS138, 2001, and derived strains.

Near the left tip of chromosome K, ten essential genes in a row contained numerous transposon insertions in the 2001u pools but not the BG2u pools. This finding suggested the left tip of chromosome K may be duplicated in the 2001u strain, causing all essential genes in the region to become non-essential. Several previous studies support this hypothesis. First, pulse-field gel electrophoresis revealed approximately 130 kb larger size of chromosome K in 2001-related strains but not in its parent strain CBS138 (Bader et al. 2012). Second, short-read sequencing of a 2001-related strain revealed approximately 2-fold increased number of reads mapping to the 130 kb tip of chromosome K relative to all other parts of the genome (Brunke et al. 2014). Third, the same region contained approximately 2-fold more *piggyBac* transposon insertions relative to other parts of the genome (Chow et al. 2024). To determine the structure of this tandem duplication, our Tn-seq reads were filtered for instances when the mate-pair read mapped much farther than the ∼400 bp distance typically observed in our sequencing libraries. Assembly of these non-contiguous reads revealed a novel DNA junction on chromosome K, where nucleotide 133,777 was adjacent to nucleotide 2,675 in the same orientation (coordinates from the CBS138 v2 reference genome (Xu et al. 2020)). These findings suggest a tandem duplication of 131,103 bp of chromosome K in 2001u that was not present in the CBS138 reference strain. To validate the tandem duplication, PCR primer sets were designed to amplify both the novel junction and the original junction (see Methods). The parental junction (1152 bp product) was observed in BG2, CBS138, and 2001 strains while the novel junction (794 bp product) was amplified only from 2001 and derived strains such as 2001HTL, the parent of hundreds of individual gene knockout mutants (Schwarzmuller et al. 2014). Two additional pools of *Hermes* insertion mutants were generated in the authenticated CBS138u strain. Tn-seq of these pools revealed almost no insertions within the 10 essential genes on the left tip of chromosome K similar to the BG2u pools and different from the 2001u pools containing the duplication. We conclude that these 10 genes are all essential in haploid strains but become non-essential when the chromosomal segment is tandemly duplicated in 2001 and derived strains.

### Transposon insertion biases reveal a paracentric inversion on chromosome L and heterochromatin deficiencies in most subtelomeric regions

The tip of chromosome K illustrated another visible difference in transposon insertion patterns: much greater insertion densities in the subtelomeres of both CBS138u and 2001u relative to BG2u insertion densities obtained previously (Gale et al. 2023) (Fig. 1A). To quantify this effect genome-wide, the insertions within a sliding window of 500 adjacent sites were totaled, normalized, and ratioed to the total insertions within the same segment of BG2u. When charted after log-2 transformation, 18 of the 26 subtelomeres exhibited 8- to more than 1000-fold increased transposon in both CBS138u and 2001u relative to BG2u, including the chromosome K subtelomere (Fig. 2). The duplicated tip of chromosome K in 2001u exhibited roughly 2-fold more insertions than the non-duplicated segment in CBS138 (Fig. 2).

**Figure 2.**
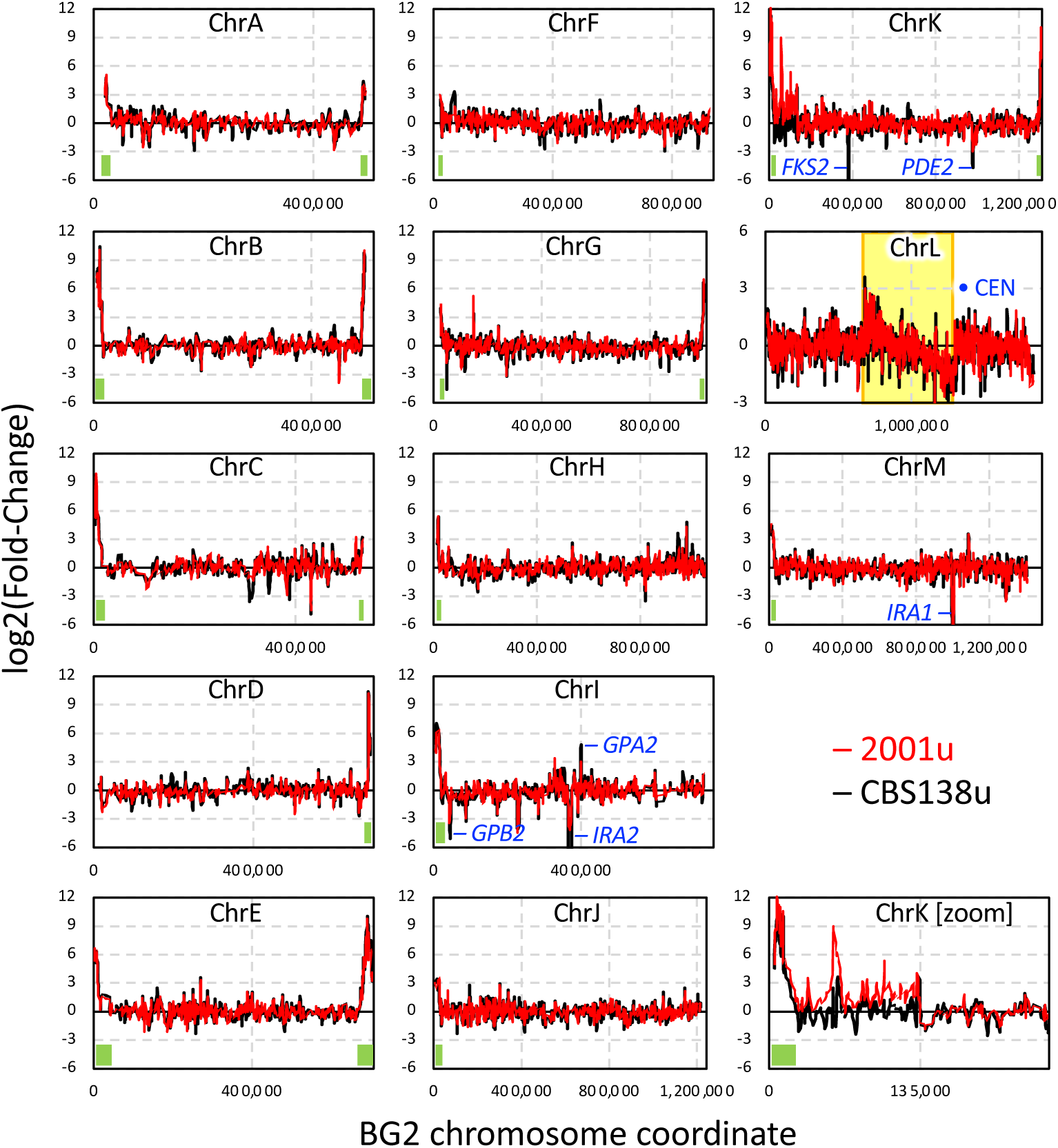
Sliding window analysis of transposon insertion densities in CBS138u and 2001u strains relative to BG2u strains. Transposon insertions at all sites in CBS138u (red) and 2001u (black) were ratioed to those of BG2u in 500-site sliding windows across all thirteen chromosomes (charted separately after taking base-2 logarithms). Eighteen subtelomeric segments with > 8-fold insertion densities are marked (green bars). On chromosome L, the approximate positions of a 632 kb inversion (yellow) and the centromere (blue dot) are illustrated. A magnified view of the left arm of chromosome K is shown separately (zoom).

The large differences in transposon accessibility at subtelomeres cannot be explained by copy-number variations or fitness differences. Rather, differences in chromatin states are likely, which is supported by several previous observations. Expression of subtelomeric genes (adhesins) have been shown to be much higher in CBS138 strains than in BG2 strains (Halliwell et al. 2012; Vale-Silva et al. 2016). CBS138 strains express a variant of Sir3, part of the SIR complex that establishes heterochromatin and silences the expression of subtelomeric adhesin genes (De Las Penas et al. 2003; Orta-Zavalza et al. 2013), containing multiple amino-acid substitutions that have been shown to disrupt subtelomeric gene silencing (Leiva-Pelaez et al. 2018; Martinez-Jimenez et al. 2013). Because transposon accessibility to DNA is highly sensitive to chromatin compaction (Gangadharan et al. 2010; Levitan et al. 2020), our Tn-seq findings suggest CBS138 strains may dysregulate expression of adhesin genes at most subtelomeres through an inability to maintain compacted heterochromatin.

Relative to BG2, CBS138 also contains a 631.6 kb paracentric inversion on chromosome L (nucleotides 679,701– 1,311,277) ending only 47.6 kb from the centromere. As *Hermes* insertions are mildly biased towards centromeres through local hopping (Gale et al. 2020), the inverted segment of chromosome L exhibits the expected gradient of transposon insertion density in CBS138u and 2001u relative to BG14 (Fig. 1B, yellow shading). These findings show that Tn-seq can detect certain types of structural variation (paracentric inversions, duplications) and chromatin alterations that were not detected using the *piggyBac* transposon in derivatives of 2001u (Chow et al. 2024).

### Fitness differences between orthologous genes in CBS138, 2001, and BG2

Several small segments of the CBS138u and 2001u genomes were visibly underrepresented with transposons relative to the BG2u genome (Fig. 2). Four of these segments spanned genes (*IRA1*, *IRA2*, *PDE2*, *GPB2*) that are all known to down-regulate protein kinase A (PKA) signaling in other yeasts (Caza and Kronstad 2019). Conversely, that up-regulates PKA signaling (*GPA2*) was visibly overrepresented with transposon insertions (Fig. 2). To quantify the significance of these outliers, transposon insertions were tabulated for all annotated genes individually and Z-scores were calculated gene-wise for CBS138u and 2001u relative to BG2u (Table S1). All five genes were significant (|Z| > 2.0) in both strains. Likewise, ten additional genes that function in the same regulatory network (*GPR1*, *RGS2, CDC25, GPB1*, *PDE2*, *YGK3*, *MSN4*, *SNF1*, *MIG1*, *NRG1*) were also found to be significant in one or both strains (Fig. 3A). However, one of these genes (*NRG1*) was discounted because it lies within a 15 kb segment of extreme sequence divergence between CBS138u and BG2u where the vast majority of 75-mers from the CBS138 reference genome fail to map uniquely to the BG2 reference genome using standard cutoffs (Fig. 3B). When the transposon insertions were remapped to the CBS138 reference genome, this 15 kb segment including *NRG1* contained numerous mappable insertions (Fig. 3B, bottom tracks) similar to the transposon insertions of BG2u mapped to the BG2 reference genome. These findings suggest that BG2 and CBS138 parent strains may differ in glucose sensing and utilization. To test this , the *IRA1* gene was knocked out in BG2u, CBS138u, and 2001u strains and competitive growth was measured in SCD medium containing 2% glucose. The *ira1Δ* mutant did not alter competitive growth in the BG2 background but did lower competitive growth in both the CBS138 and 2001 strain backgrounds (Fig. 3C). These findings validate the transposon insertion findings for *IRA1* and suggest that the glucose sensing pathway differentially alters fitness of BG2 and CBS138 parent strains. BG2 and CBS138 were shown previously to consume glucose at different rates during exponential phase growth and to achieve different cell densities in stationary phase (Legrand et al. 2016). These effects may be due in part to large sequence variation within *NRG1* coding sequences (only 85.7% amino acid identity between CBS138 and BG2), which encodes a glucose-sensitive transcription repressor in *S. cerevisiae* (Lee et al. 2013).

**Figure 3.**
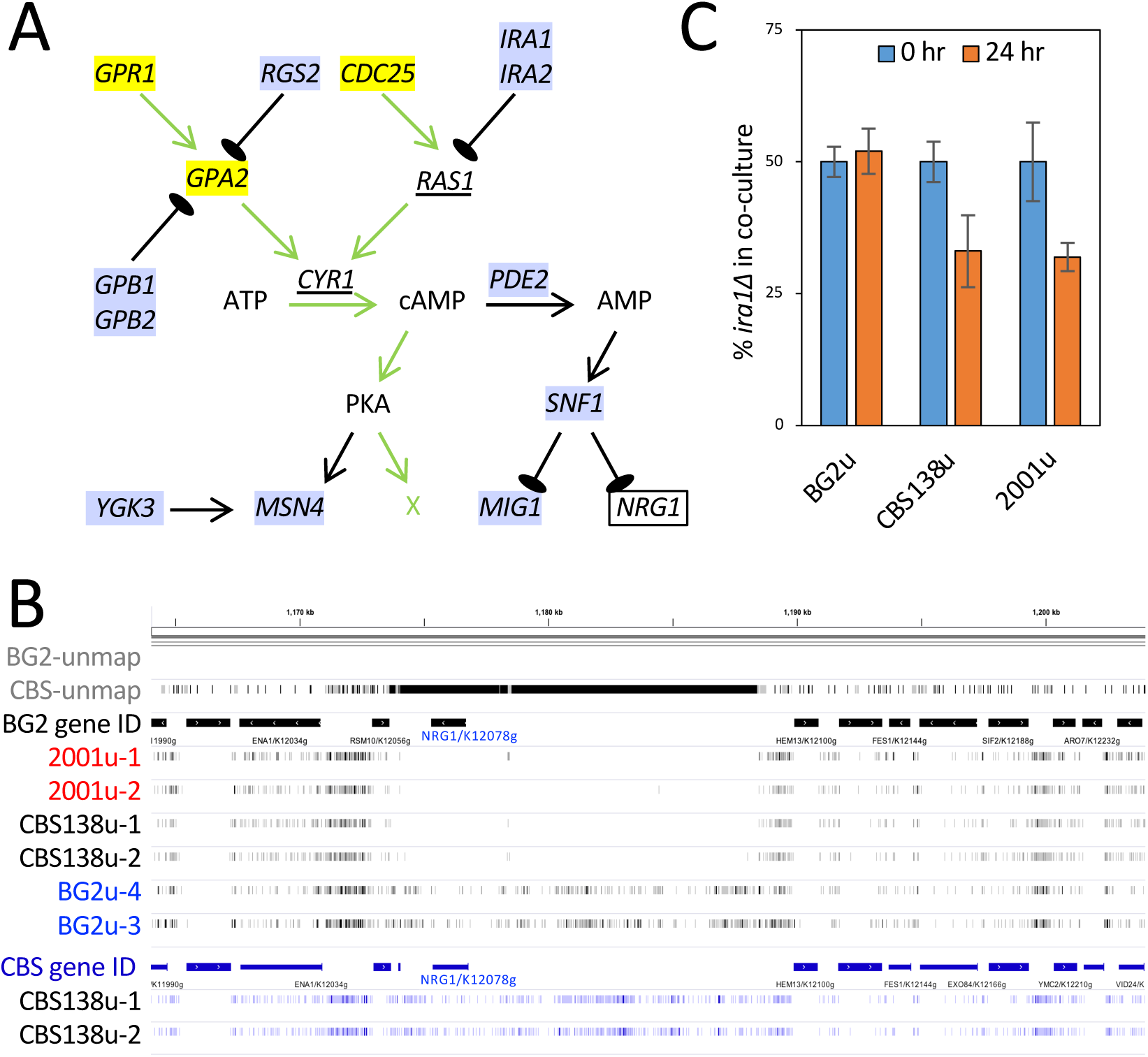
Fitness effects of insertions in genes that sense glucose. (A) Model of the glucose sensing gene products of *S. cerevisiae* that are differentially disrupted by transposons in strains CBS138u and 2001u relative to BG2u. Yellow highlighted gene names are significantly over-represented in CBS138 and 2001u relative to BG2u whereas significantly under-represented genes are highlighted in blue. Essential genes are underscored. Unmappable genes are boxed. Genes encoding protein kinase A (PKA), which responds to cyclic-AMP (cAMP) are not significant and not shown. Positive and negative interactions are indicated with pointed and blunt arrows, respectively. Green arrows indicated the path required for cell growth through partially unknown targets (X). (B) IGV browser view of a 40 kb segment of chromosome K surrounding the *NRG1* gene. The top tracks (black) utilized the BG2 reference genome while the bottom tracks (purple) utilized the CBS138 reference genome. (C) Several *ira1Δ* knockout mutants and their parent strains were mixed at equal ratios, diluted, co-cultivated for 24 hr in SCD medium, plated for single colonies, then replica plated to SCD-ura medium to quantify percentages of the *ira1Δ* strain in the mixed cultures. Columns represent the averages (±SD) of four biological replicates.

After excluding subtelomeric, inverted, and duplicated segments, a total of 43 genes exhibited significantly lower Z-scores while 14 genes had significantly higher Z-scores in both the CBS138u and 2001u strains relative to BG2u. Three genes with high Z-scores (*YPD1*, *SLN1*, *SSK2*) function together in an osmo-sensing pathway in BG2 that has been shown to be partially defective in CBS138-derived strains (Gregori et al. 2007). Genes with low Z-scores represented several functional categories. The most striking category involved the N-glycosylation process in the endoplasmic reticulum in which four non-essential genes (*ALG3, ALG5, ALG6, ALG8*) were strongly under-represented with transposon insertions. Knockout mutants of *ALG3*, *ALG5* or *ALG6* all resulted in a lower growth rate at elevated temperature in a 2001u-related background (Schwarzmuller et al. 2014) but not in the BG2u background (Fig. S1). Another 87 and 43 genes exhibited significantly lower and higher Z-scores respectively in only the CBS138u strain but not the 2001u strain. Those numbers are much larger than the additional 21 and 12 genes that exhibited significantly lower and higher Z-scores in the 2001u strain but not CBS138u. This difference raises the possibility that the tandem duplication on chromosome K confers a more BG2-like fitness landscape to the CBS138 strain background without significantly altering the states of subtelomeric chromatin.

To focus more directly on the fitness effects of the 131 kb tandem duplication, the sequenced transposon insertions in 2001u, CBS138u, and BG2u were re-mapped to the CBS138 reference genome instead of BG2 used in earlier sections, and then Z-scores were calculated for all genes between the two strains (Table S2). Most of the genes within the duplicated segment (42 of 60) were significantly over-represented with transposon insertions in 2001u relative to CBS138 while most of the remainder (15 of 18) were marginally over-represented. Interestingly, most genes in the glucose sensing pathway (Fig. 3A) were significantly overrepresented in these 2001-to-CBS138 comparisons but underrepresented in the CBS138-to-BG2 comparisons (Table S1). The divergence of CBS138u fitness profiles from both 2001u and BG2u again suggests that the tandem duplication on chromosome K may convert the physiology of CBS138 to a state that more closely resembles that of BG2.

Dozens of other genes were significantly over-represented with transposon insertions in 2001u relative to CBS138, including several genes involved in protein O-glycosylation (*PMT2*, *KRE2*, *MNN2*, *VAN1*). The most striking example of differential representation with transposon insertions was the *FKS2* gene, which encodes a beta-1,3-glucan synthase in the cell membrane (Katiyar et al. 2012). Insertions within *FKS2* were highly elevated in 2001u relative to CBS138u (Z = 8.1) where the gene seemed nearly essential (Fig. 4). The difference was easily visualized by scanning window analysis (Fig. 2, ChrK). Smaller, but still significant, effects were seen for *FKS1* (Z = 2.6), which encodes a second beta-1,3-glucan synthase that is functionally redundant with *FKS2* (Katiyar et al. 2012). As insertions in these genes are unlikely to improve fitness of any strain, they likely have a greater fitness defect in CBS138u than in 2001u and BG2u. Consistent with this hypothesis, insertions in *LRG1*, which encodes a negative regulator of the beta-1,3-glucan synthases (Watanabe et al. 2001), were significantly over-represented in CBS138u relative to 2001u and BG2u (Z = 4.3 and 2.5, respectively). Several remodelers of cell wall beta-1,3-glucan in the cell wall (*SCW4*, *SCW10*, *GAS1-B*) also exhibited significant differences between CBS138u and 2001u (Z = 4.7, -2.7, 4.2) and not BG2u. Altogether, these findings suggest that synthesis and remodeling of beta-1,3-glucan in the cell wall may be altered in CBS138 relative to BG2 and that the tandem duplication on chromosome K in 2001 partially reverses the effect.

**Figure 4.**
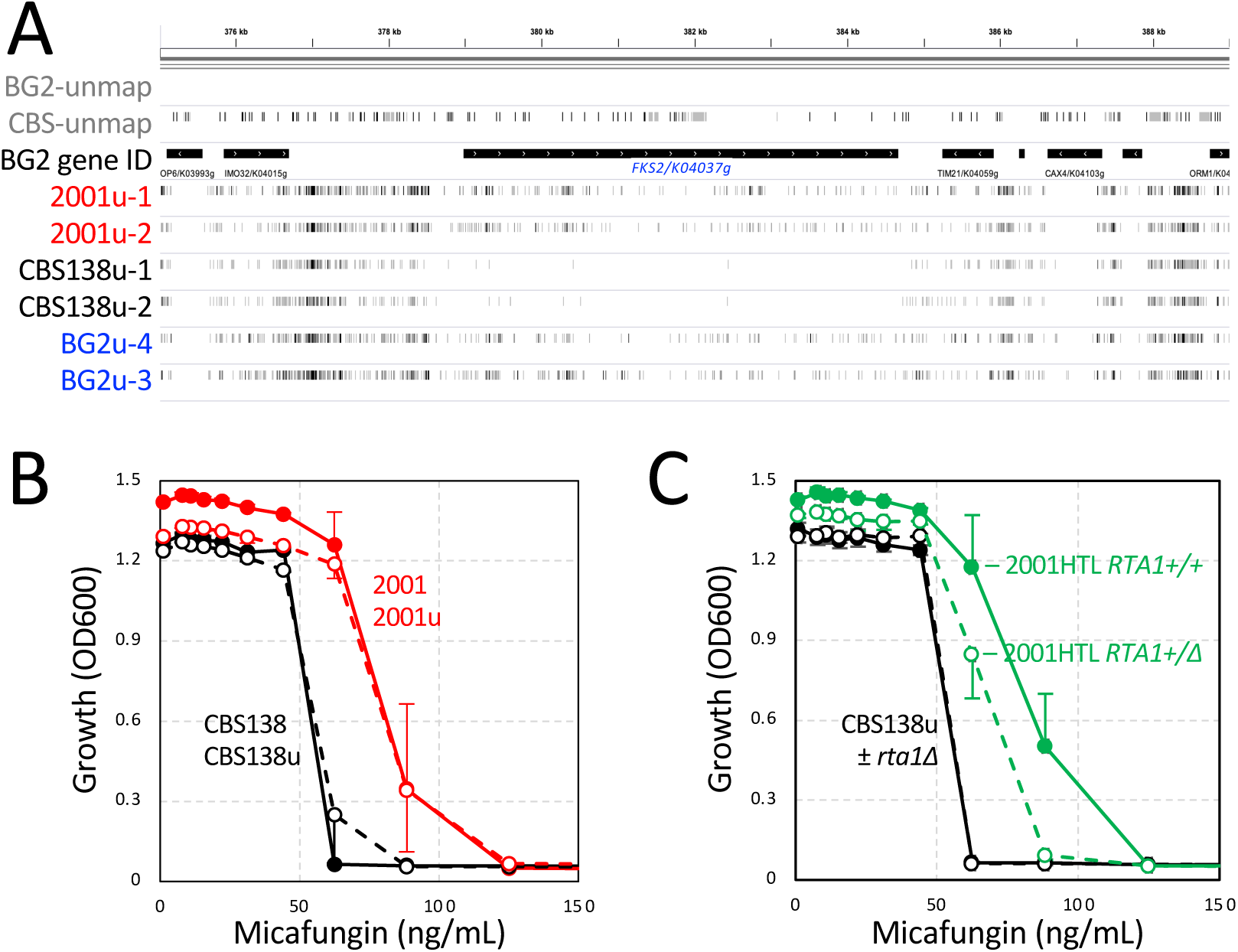
Phenotypic variation involving *FKS2* and micafungin. (A) IGV browser view of transposon insertions near the *FKS2* gene locus. (B, C) Growth of the indicated strains was measured after 48 hr incubation in SCD medium containing varying doses of micafungin. Symbols indicate the averages of four (B) or three (C) biological replicates (±SD).

To test this, the resistances of CBS138u and 2001u to micafungin, an echinocandin-class inhibitor of the *FKS1* and *FKS2* gene products (Eschenauer et al. 2007), were quantitatively assessed. The 50% inhibitory concentration (IC50) of micafungin was consistently 1.5-fold higher in 2001u than CBS138u (Fig. S2). Similar results were obtained previously using caspofungin, another echinocandin-class inhibitor of beta-1,3-glucan synthases, and using other types of cell wall stressors (Bader et al. 2012). These findings suggest that increased dosage of genes within the 60-gene tandem duplication confers several phenotypes such as resistance to echinocandins, tolerance to *FKS2* insertions, altered cell wall biogenesis, and altered glucose sensing, which often revert CBS138 to a more BG2-like state in laboratory conditions.

### Genes contributing to micafungin resistance

To explore the genetic basis of micafungin resistance, pools of transposon insertion mutants in both CBS138u and 2001u strains were grown to saturation, diluted 100-fold into fresh medium containing varying concentrations of micafungin, and shaken for 48 hours. At the lowest doses, the cultures reached stationary phase by 16 hours whereas 48 hours was necessary to reach stationary phase at the highest doses (Fig. S3). The insertion sites in each culture were then amplified, sequenced, mapped to the CBS138 reference genome, and tabulated gene-wise as before. Additionally, a second gene-wise tabulation was performed where the most frequently sequenced insertion site in each gene was identified and the sequence reads at that site were subtracted from the total count for that gene. This “top-site removed” (TSR) tabulation was implemented to flag potential passenger effects (Michel et al. 2017), where fitness advantages are conferred by spontaneous mutations in the genome rather than the transposon insertion itself (for example, gain-of-function point mutations within the *FKS1* or *FKS2* genes that diminish micafungin binding and/or increase enzymatic activity (Garcia-Effron et al. 2009; Park et al. 2005)). Z-scores were then calculated from the tabulated data for each gene at every dose of micafungin relative to the untreated control (Table S2). At low and moderate doses of micafungin, few genes were affected by the TSR approach. However, at the highest micafungin doses where selective pressures were largest, dozens of genes exhibited strongly positive Z-scores that were neutralized by omitting the single top-site from the tabulations (Table S2). The TSR-sensitive genes in the CBS138u datasets generally did not overlap with those of the 2001u datasets, as expected from random passenger effects. In both strain backgrounds with and without the TSR modification, the *FKS1* gene was significantly under-represented with transposon insertions relative to untreated control (Z-scores < -2.7) at nearly all doses of micafungin (Table S2). These findings confirm previous studies where *ffis1Δ* knockout mutants in the 2001 background exhibited lower resistance to micafungin (Katiyar et al. 2012) and confirm the experimental conditions were suitable for identifying additional regulators of micafungin resistance.

Figure 5A shows the Z-scores obtained from 2001u (16 ng/mL micafungin) charted with those of CBS138u (8 ng/mL micafungin) after the TSR tabulation to remove most passenger effects. These doses were chosen based on a similar level of growth inhibition during micafungin treatment (Fig. S3). Overall, the two independent datasets obtained at moderate micafungin doses were remarkably well correlated (PCC = 0.91). Of the 60 genes located within the tandem duplication on chromosome K, only a few (*RTA1, SAC1, DSC2*) appeared significantly less resistant to micafungin in the 2001u strain but not the CBS138u strain. In accordance with this, hemizygous *rta1Δ/RTA1* knockout mutants exhibited slightly lower resistance to micafungin relative to the control strain 2001HTL whereas complete *rta1Δ* knockout mutants did not exhibit lower micafungin resistance in CBS138u background (Fig. 4C). These findings suggest that duplication of *RTA1* in 2001-derived strains confers partial resistance to micafungin while other genes within the duplicated segment confer additional resistance. Three genes within the tandem duplication (*MRPL25*, *MRP10*, *MTF2*) produced very high Z-scores in CBS138u but not in 2001u (Fig. 5A), suggesting that insertions within one allele are fully complemented by the remaining allele in 2001u. Interestingly, all three of the genes encode mitochondrial proteins. Outside of the duplication, another 184 genes that all encode mitochondrial proteins exhibited very high Z-scores in CBS138u and/or 2001u (Fig. 5A, yellow circles). In BG2u, insertions in the same mitochondrial genes conferred high resistance to fluconazole (Gale et al. 2020). When Z-scores of CBS138u exposed to micafungin were plotted against Z-scores of BG2u exposed to fluconazole, a good correlation was observed (PCC = 0.68; Fig. 5B). Together, these findings from different strains and different classes of antifungals suggest that mitochondrial dysfunction can confer pleiotropic drug resistance.

**Figure 5.**
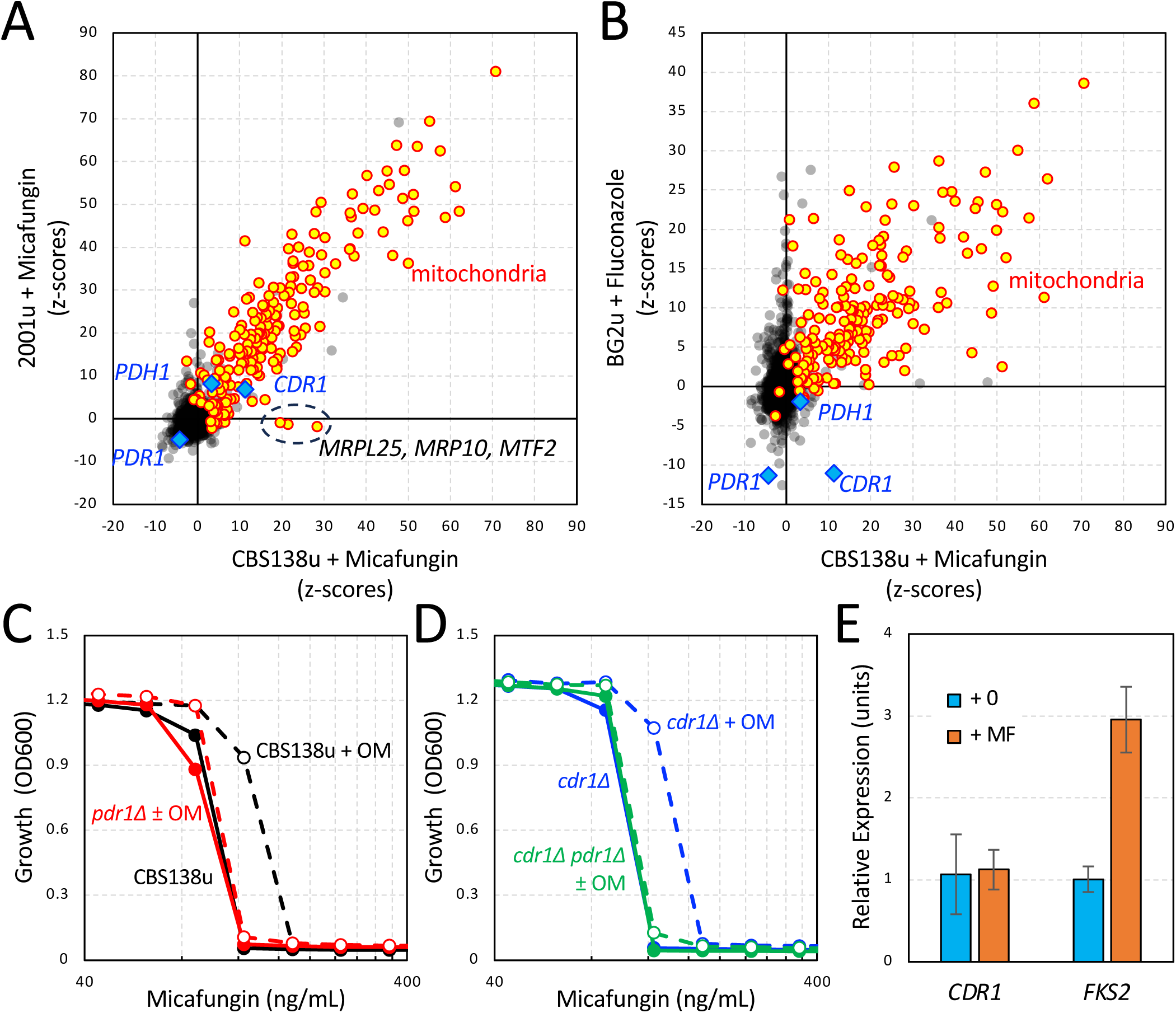
Identification of genes that regulate resistance to micafungin. Z-scores of all annotated genes in CBS138 were calculated for tabulated transposon insertions in CBS138u exposed to 8 ng/mL micafungin and charted against Z-scores calculated similarly for 2001u exposed to 16 ng/mL micafungin (A) and for BG2u exposed to 128 µg/mL fluconazole (B). Significant mitochondrial genes (yellow) and *PDR1*-related genes (blue) were layered onto charts of all other genes (gray, black). Selected genes were annotated. (C, D) Micafungin resistance assays were performed on the indicated strains after growth for 20 hr in SCD medium containing (dashed lines) or lacking (solid lines) 40 µg/mL oligomycin. (E) mRNA levels derived from the *CDR1* and *FKS2* genes relative to *PGK1* (control) were quantified by qRT-PCR after 1 hr exposure of strain CBS138u to 0.12 µg/mL micafungin. Columns show averages of three biological replicates (±SD).

Mitochondrial dysfunction strongly activates the Pdr1 transcription factor in *C. glabrata*, which induces expression of *CDR1* and other target genes encoding ABC transporters that together promote pleiotropic drug resistance by exporting diverse antifungals (Chow et al. 2024; Gale et al. 2023). Consistent with a role for Pdr1 in micafungin resistance, *PDR1* was significantly under-represented with transposon insertions in both CBS138u and 2001u (Z = -4.4, -5.2) after exposure to micafungin (8 ng/mL, 16 ng/mL), similar to fluconazole exposure in BG2u (Z = -11 (Gale et al. 2020)). Surprisingly, *CDR1* was over-represented with transposon insertions in CBS138u and 2001u during micafungin exposure (Z = +12, +8.7), which is opposite to the effects of *CDR1* on fluconazole resistance (Z = -11). A similar, but less pronounced effect was observed for a paralog of *CDR1*, termed *PDH1*, whose expression is also induced by activated Pdr1 (Z = +3.6, +7.9). These findings suggest that Pdr1 may increase micafungin resistance in *C. glabrata* through a complex process where some of its targets may increase resistance and others (e.g. *CDR1*, *PDH1*) may decrease resistance.

These findings predict that oligomycin, an inhibitor of mitochondrial F1F0-ATPase that strongly activates Pdr1 (Gale et al. 2023), may increase micafungin resistance in a *PDR1-*dependent and *CDR1*-independent fashion. To test these predictions, micafungin resistance was quantified for *pdr1Δ*, *cdr1Δ*, and double knockout mutants in the CBS138u strain background in both the presence and absence of oligomycin. Oligomycin significantly increased resistance to micafungin in CBS138u but not in *pdr1Δ* mutants (Fig. 5C). Similarly, oligomycin increased micafungin resistance in *cdr1Δ* mutants but not in *cdr1Δ pdr1Δ* double knockout mutants (Fig. 5D). In the absence of oligomycin, *pdr1Δ* mutants were slightly less resistant to micafungin while *cdr1Δ* mutants were slightly more resistant to oligomycin. These findings confirm the significant Z-scores obtained from the Tn-seq datasets and confirm the prediction that activated Pdr1 can promote micafungin resistance through *CDR1*-independent effects.

To determine whether micafungin exposure can activate Pdr1, *CDR1* mRNA levels were quantified by qRT-PCR after 1 hour exposure to micafungin. The CBS138u strain exhibited no significant increase in *CDR1* expression after exposure to micafungin (Fig. 5E). In contrast, micafungin exposure strongly induced expression of *FKS2* mRNA, which depends on calcineurin and Crz1 signaling (Pavesic et al. 2023). These data suggest that micafungin exposure does not readily activate Pdr1 in these conditions or that *CDR1* expression somehow becomes insensitive to activated Pdr1. Identification of Pdr1 targets that promote resistance to micafungin could help reveal the underlying mechanism.

### Other determinants of micafungin resistance

Genes that innately increase resistance to micafungin represent new targets for development of echinocandin-enhancing drugs. To identify such genes, the Z-scores from CBS138u and 2001u were averaged and sorted. At the lowest effective doses of micafungin (2 µg/mL for CBS138u and 4 µg/mL for 2001u), *FKS1* (|Z| = -4.1) and six additional genes exhibited average Z-scores less than -3.0 (Table S2). Two of these genes (*LAF1*, *IPT1*) have been identified as targets of Pdr1 (Paul et al. 2014; Vermitsky et al. 2006) and therefore candidates for Pdr1-dependent micafungin resistance. Two other genes (*MID2*, *YND1*) play roles in cell wall biosynthesis and were found previously to confer resistance to fluconazole in strain BG2u (Z = -6.1, -7.2 (Gale et al. 2020)). *MID2* encodes a transmembrane sensor of cell wall damage that compensates for cell wall deficiencies by activating the small GTPase Rho1, which activates the beta-1,3-glucan synthases and regulates other targets (Sekiya-Kawasaki et al. 2002). *YND1* encodes a Golgi-localized apyrase that promotes biosynthesis of mannan and glucan components of the cell wall by converting luminal GDP and UDP to mononucleotides that are then exchanged for the cytoplasmic nucleotide-sugars GDP-mannose and UDP-glucose (Gao et al. 1999). At higher doses of micafungin (4 µg/mL for CBS138u and 8 µg/mL for 2001u) another 43 genes exhibited average Z-scores less than -3.0, 4 of which encode mannosyltransferases of the Golgi that depend on GDP-mannose import for biosynthesis of cell wall mannan (*MNN4*, *KTR6*, *MNN5*, *HOC1*). Several genes involved in chitin biosynthesis (*CHS3*, *PFA4*) and lipid homeostasis (*IZH3*, *ARE2*, *OSH3*, *YSP2*, *OSH6*, *DNF1*) were also significant in these conditions. Two genes involved in N-glycan biosynthesis (*ALG12*) and sphingolipid biosynthesis (*LAG1*) in the ER were also recovered. All of these non-essential genes could be targeted for development of drugs that might enhance efficacy of micafungin.

Drugging of essential genes in the same pathways as the non-essential genes described above may be even more effective as echinocandin co-drugs, as they would also inhibit growth of *C. glabrata* cells when used alone. To test this concept, we determined whether the inhibitory effects of micafungin can be enhanced by SDZ 90-215 and myriocin. SDZ 90-215 blocks an essential nucleotide-sugar exchanger in the Golgi encoded by *VRG4* whose function is required for mannosyltransferase activity and biosynthesis of mannan (Snyder et al. 2019). Myriocin blocks an essential serine-palmitoyltransferase in the ER, which is encoded by *LCB1* and *LCB2*, that synthesizes a lipid precursor of sphingolipids (Liu et al. 2013). Titrating concentrations of these compounds were mixed with micafungin titrations and the cocktails were tested for their ability to inhibit growth of CBS138 and CBS138u strains in a checkerboard assay. After 24 hr incubation in SCD medium, cell growth was measured and analyzed quantitatively to determine the IC50 of each compound alone and the combination index (CI), or the IC50 of the two compounds combined in 1:1 ratio (Snyder et al. 2019). SDZ 90-215 and micafungin exhibited additive interactions in both strains (CI ∼ 1.0; Fig. 6). Myriocin and micafungin exhibited slightly synergistic interactions in both strains (CI < 1.0; Fig. 6). In contrast, manogepix, an experimental antifungal that blocks GPI anchor biosynthesis in the ER (Kapoor et al. 2019; Miyazaki et al. 2011; Watanabe et al. 2012), and micafungin did not exhibit additivity or synergism and instead seemed independent (CI ∼ 2.0). None of these combinations exhibited antagonism (CI > 2.0). These findings confirm the concept that unbiased genome-wide screening can yield new targets for enhancing the efficacy of existing antifungals.

**Figure 6.**
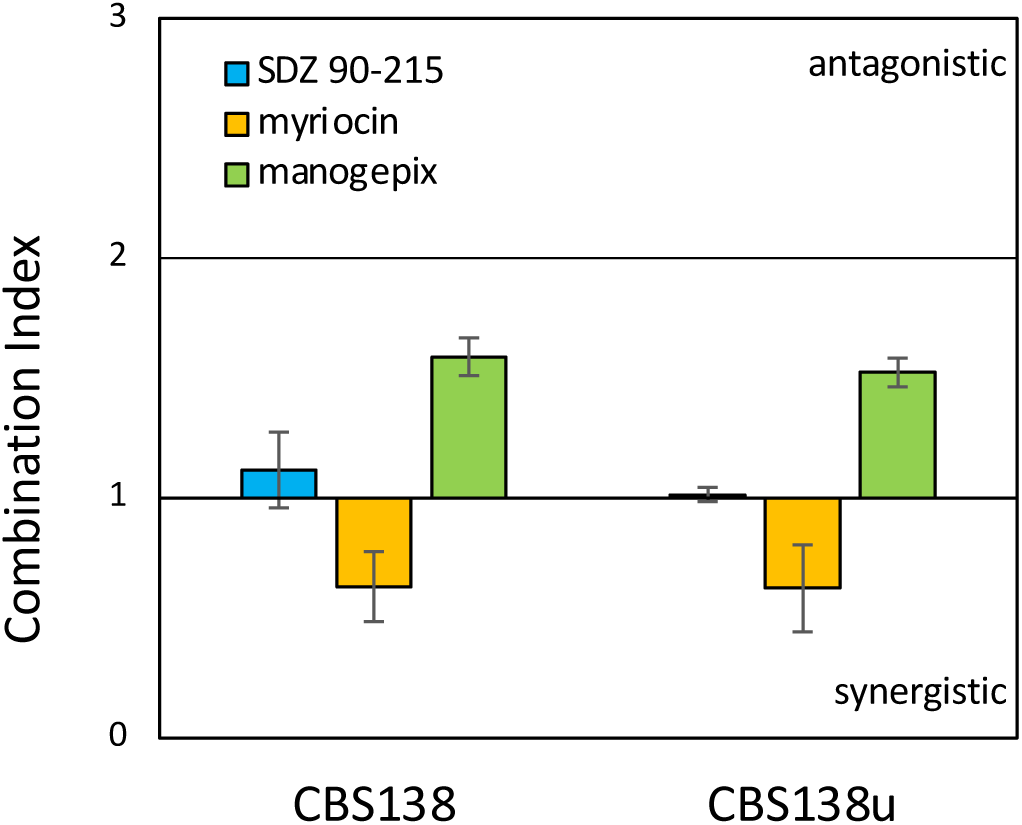
Synergy between micafungin and experimental antifungals. Checkerboard assays were performed where micafungin concentration was co-varied with an experimental antifungal in SCD medium and growth of CBS138 or CBS138u strains was measured after 24 hr incubation. Combination Index was then calculated for each checkerboard (columns indicate averages of 2 to 4 replicates, ±SD).

## DISCUSSION

This study comprehensively probes the genome of the *C. glabrata* reference strain CBS138 with transposon insertional mutagenesis for the first time. In comparison to strain BG2, which is nearly 0.9% different at the nucleotide level (Xu et al. 2021), CBS138 exhibited dramatically increased transposon insertions in the subtelomeres of most chromosomes. The increased transposon accessibility likely reflects a more open state of chromatin in those subtelomeres. This is consistent with previous studies that showed elevated expression of subtelomeric adhesin genes and increased adhesion of CBS138-related strains relative to BG2 and other strains (Halliwell et al. 2012; Vale-Silva et al. 2016). Non-synonymous mutations in the *SIR3* gene of CBS138 that alter sirtuin function at subtelomeres have been proposed as a cause of chromatin decompaction and adhesin expression (Leiva-Pelaez et al. 2018; Martinez-Jimenez et al. 2013). With over 12,500 non-synonymous mutations between CBS138 and BG2 (Xu et al. 2021), additional variants may also contribute to the observed heterogeneity in chromatin and produce an array of phenotypic differences within the strains. Examples include altered fitness effects of disrupting genes involved in glucose sensing (Fig. 2), osmolarity sensing (*YPD1, SLN1, SSK2*), and cell wall biogenesis (*FKS1*). With nearly 1% sequence diversity between the major clades of *C. glabrata* strains (Carrete et al. 2018; Marcet-Houben et al. 2022), large phenotypic diversity is likely to exist. The portability of transposon mutagenesis approaches *in vivo* will facilitate studies of this phenotypic diversity.

A derivative of CBS138 known as 2001 was found here to contain a 131 kb tandem duplication of the left tip of chromosome K, in concordance with previous observations of increased chromosome length (Bader et al. 2012) and increased sequencing coverage (Brunke et al. 2014). This duplication spans 60 genes, 10 of which were classified essential in BG2 (Gale et al. 2020) and also appeared essential in CBS138. Strain 2001 has been used to generate the 2001HTL and KUE100 strains employed in large-scale gene knockout projects (Schwarzmuller et al. 2014; Ueno et al. 2007). Recently, the 2001HTL strain was mutagenized with the *piggyBac* transposon and analyzed by Tn-seq (Chow et al. 2024). Few studies with 2001-derived strains report the existence of the tandem duplication and many studies conflate this strain with authentic CBS138. However, numerous phenotypic differences may exist between 2001 and CBS138. In addition to cell wall differences reported previously (Bader et al. 2012), we observed different fitness consequences for insertions in dozens of genes, particularly those involved in glucose sensing and glucan biosynthesis. In these cases, the fitness effects of insertions in 2001u more closely resembled those of BG2u than those of CBS138u in the laboratory conditions employed here. The tandem duplication in 2001 may therefore improve fitness of CBS138 in laboratory conditions. CBS138 grows to half the density achieved by BG2 in glucose medium (Legrand et al. 2016) and is very closely related to a rare environmental isolate of *C. glabrata* known as M202019 (Xu et al. 2016). CBS138 may have been recently acquired as a pathogen of humans and somewhat atypical representative of the *C. glabrata* species complex. More work on the diversity and phylogenetics of *C. glabrata* isolates could provide a more comprehensive picture of this pathogen as well as the best ways of controlling it.

A whole-genome approach to the genetic basis of micafungin resistance *in vitro* was also implemented for the first time in both CBS138- and 2001-derived strains. Though 2001 was slightly more resistant to this echinocandin and others (Bader et al. 2012), the genes regulating micafungin resistance *in vitro* were remarkably consistent in both strains (Fig. 5A). Surprisingly, the *PDR1* gene was important for competitive fitness at low concentrations of micafungin, and transposon insertions that eliminate its negative regulators (e.g. mitochondrial functioning) strongly increased relative fitness in micafungin. Oligomycin, which generates mitochondrial stresses and activates the Pdr1 transcription factor (Gale et al. 2023), increased resistance to micafungin when added to CBS138u cells but not to *pdr1Δ* cells. This effect was independent of *CDR1*, which encodes an ABC transporter involved in efflux of fluconazole and cycloheximide (Vermitsky et al. 2006). Oligomycin and Pdr1 therefore conferred micafungin resistance through other target(s). One target of Pdr1 that contributed to micafungin resistance in this study is *IPT1*, which promotes biosynthesis of a sphingolipid (Shahi et al. 2022). On the other hand, *CDR1* and its paralog *PDH1* may have the opposite effect as transposon insertions in these genes increased competitive fitness in micafungin. The antagonistic effects of the different Pdr1 targets and the inability of micafungin stress to activate Pdr1 (Fig. 5E) complicates the picture of how this stress-sensitive transcription factor contributes to echinocandin resistance. A key missing element of the regulatory network may be calcineurin, which was recently shown to bind and regulate Pdr1 activity in *C. glabrata* (Vu et al. 2023).

The Tn-seq screens performed here reveal additional genes that contribute to fitness during micafungin exposure in *C. glabrata* and provide new insights into improving echinocandin efficacy. Dozens of genes in both the CBS138u and 2001u strain backgrounds exhibited significantly negative Z-scores (diminished relative fitness) at low doses of micafungin. All of these genes are non-essential and therefore unlikely to become druggable targets for improving micafungin efficacy. Essential genes that operate in the same pathways as these non-essential hits are more promising. As several different mannosyltransferases of the Golgi complex were significant, we explored the possibility that SDZ 90-215, an inhibitor of Vrg4 in the Golgi that supplies the mannosyltransferases with critical substrates (GDP-mannose) (Snyder et al. 2019), can increase the potency of micafungin *in vitro*. In checkerboard assays, SDZ 90-215 and micafungin behaved in an additive fashion when combined at equal dosages (Fig. 6). Myriocin and micafungin combinations were tested similarly and found to interact even more strongly (CI = 0.63) while manogepix and tunicamycin exhibited no interaction (CI = 1.55). Myriocin blocks essential enzymes in the ER required for sphingolipid biosynthesis while manogepix potently blocks enzymes in the ER required for GPI anchor biosynthesis and is currently being developed as novel broad-spectrum antifungal (Hodges et al. 2023; Pappas et al. 2023). While these findings are promising, more research will be needed to establish additivity and synergy of the compounds in multiple *C. glabrata* strains and in multiple environments, including infected host animals.

The Tn-seq approach has recently enabled genome-wide forward genetic screens in eukaryotic pathogens such as *C. glabrata* (Gale et al. 2020), *C. albicans* (Segal et al. 2018), *C. auris* (Gao et al. 2021), and *Plasmodium falciparum* (Zhang et al. 2018). Common goals in all these systems are to generate pools of insertion mutants that contain high complexity, low insertion biases, and easily interpretable datasets obtained after exposing the pools to different stresses and environments. The *miniDS* transposon exhibits no sequence preference at insertion sites but can produce gain-of-function effects in some instances (Michel et al. 2017). The *piggyBac* transposon strongly prefers to insert at TTAA sites, which greatly reduces complexity of the pools (Chow et al. 2024). The *Hermes* transposon utilized here exhibits mild preference for TnnnnA sites in the genome and a modest degree of local hopping (Gale et al. 2020), producing pools of intermediate complexity while permitting identification of large chromosomal inversions such as that on chromosome L (Fig. 2). As *Hermes* insertions occurred mostly in stationary phase cultures (Gale et al. 2020), jackpot events were extremely rare. However, all these Tn-seq variations are subject to passenger effects (spontaneous mutations in the pools that confer selective advantage) which may be difficult to identify. The TSR approach implemented here flags potential passenger effects that can produce false results for individual genes. Together, the Tn-seq approaches are poised to provide numerous insights into previously intractable questions in diverse pathogens.

## METHODS

### *C. glabrata* strains and authentication

All *C. glabrata* strains and primers utilized in this study are listed in Table S3. Authentic strain CBS138 (ATCC2001) was provided by David Perlin and the derived strain 2001 (Kitada et al. 1995) containing a 131 kb duplication was obtained from Taiga Miyazaki. The CBS138u and 2001u strains containing *ura3Δ* knockout mutations were constructed by transformation of the parent strains with a template-free PCR product, selection on SCD medium containing 1 mg/mL 5-FOA, and then screening for deletion of the *URA3* locus by colony PCR. Several gene knockouts (alg5Δ, pdr1Δ, fks2Δ) were constructed by PCR amplification of the locus and selectable marker from a previously generated strain, transformation of the appropriate *C. glabrata* strain, selection on agar plates, and authentication of colonies by PCR, as indicated in Table S3. Other gene knockouts were generated using the PRODIGE method (Edlind et al. 2005), which replaces the coding sequences of the gene with coding sequences of the *S. cerevisiae* URA3 using pRS406 plasmid (Sikorski and Hieter 1989) as a template and PCR primers containing a 60-nucleotide flanking homology to the target gene. After transformation into CBS138u or 2001u, transformants were grown on SCD-uracil plates and URA+ colonies were authenticated via PCR.

To determine the nature of the chromosome K duplicated segment in 2001-derived strains, Tn-seq reads mapping near the suspected breakpoint in the CBS138 reference genome v2 (Xu et al. 2020) were compared to the mapped positions of their mate-pairs. If the difference was more than 5 kb, the reads were binned and assembled. Thousands of independent sequence reads spanned a novel DNA junction not present in the reference genome, where nucleotide 133,777 was adjacent to nucleotide 2,675 and in the same orientation (i.e. a tandem duplication of 131,102 bp). To authenticate this junction in 2001-derived strains, a simple PCR test involving three primers designed (Fig. 1B). For PCR authentication, genomic DNA was extracted by heating cells for 20 minutes at 80°C in TE-lithium acetate with 1% SDS, precipitated with 75% ethanol, then dissolved in water. PCR was performed using NEB Phusion High-Fidelity PCR Master Mix (Cat. #M0531) with 3% DMSO and a 0.6 µM concentration of each primer (F, R1, R2) listed in Table S3 using the following settings: 5-minute denaturation at 95°C, 35 cycles of 95°C for 30 seconds, 56°C for 30 seconds, and 72°C for 1 minute, and then a 7-minute final extension at 56°C. PCR products were electrophoresed on 1% agarose gels along with NEB 100bp DNA Ladder (Cat. #N3231). All 2001-derived strains contained a 794 bp band that was sequenced and found to span the novel junction (Fig. 1C).

### Transposon Mutagenesis and Tn-seq

Plasmid pCU-MET3-Hermes was transformed into strains 2001u (TJN32) and CBS138u (TJN69). Two pools of *Hermes::NAT1* insertion mutants were generated independently for each strain in SCD-ura-met-cys medium as described previously (Gale et al. 2020). Cells containing transposon insertions in the genome were enriched by growth in SCD+5FOA as described and aliquots in 15% glycerol were stored at -80°C. Genomic DNA was extracted from the pools using the Quick-DNA Fungal/Bacterial Miniprep Kit from Zymo Research (Cat. #D6005) and fragmented using a Bioruptor Pico sonication device. Frayed ends were repaired, A-tailed, and ligated to indexed splinkerette adapters using the NEBNext Ultra II DNA Library Prep Kit for Illumina (Cat. #E7645) as described previously (Gale et al. 2020). DNA was size-selected using AMPure XP magnetic beads from Beckman Coulter (Cat. #A63881) and DNA containing the HERMES transposon was amplified via PCR followed by a second PCR to add the Illumina capture sequence. These indexed libraries were sequenced using a MiSeq Reagent Kit v3 (2x75 bp; Illumina) to generate paired-end reads.

FastQ files obtained from the MiSeq were demultiplexed using CutAdapt and aligned to the BG2 v1 genome (Xu et al. 2021) or the CBS138 v2 reference genome (Xu et al. 2020) using Bowtie2 (Langmead and Salzberg 2012). Reads with Q-score => 20 and no mismatches at position 1 were retained for analysis. The number of sequence reads that map to each mapped insertion site was tabulated to create site-count files. To perform sliding window analysis of Tn-seq reads from CBS138u or 2001u relative to BG2u, a merged site-count file was generated after mapping all forward reads to the BG2 genome sequence. The datasets were normalized to matching depths and then average number of insertions per strain for each sliding window of 500 sites was calculated, incremented by 1, ratioed, converted to logarithm base-2, and charted as function of chromosomal coordinate (Fig. 2).

Insertion sites were visualized on the IGV genome browser (Robinson et al. 2011) by converting site-count files into BED tracks. Sites in the BG2 and CBS138 genomes that could not be mapped to BG2 genome sequence due to duplications, mismatches at position 1, or high sequence divergence between strains were similarly visualized. To generate the unmappable site-count files, the BG2 and CBS138 genome sequences were parsed into all possible 75-mers and then mapped to the reference genomes using Bowtie2 and the same filtering parameters as Tn-seq reads. All genomic sites that have a single mapped 75-mer were excluded and only sites with zero or >1 mapped 75-mers were included in BED files for visualization with tick marks on the IGV browser. All BED files and site-count files are available from Candida Genome Database or by request.

Frozen pools of insertion mutants were thawed, revived in SCD medium, and then diluted 100-fold from saturated cultures into SCD medium containing varying doses of Micafungin (Cayman Chemicals) as well as SCD medium alone. The cultures were shaken at 30°C, periodically sampled to determine cell density (Fig. S3), and then harvested after 48 hr. Genomic DNA was harvested from the pelleted cells and subjected to Tn-seq as described above. The sequence reads were aligned to the CBS138 reference genome to generate a new site count file, which was used to tabulate reads for all annotated genes individually. To eliminate potential passenger effects that could skew the analysis, the number of reads at the most frequently represented insertion site of each gene was subtracted from the gene-wise tabulation to generate a second top-site removed (TSR). Both tabulations were normalized and utilized to calculate Z-scores as described previously (Gale et al. 2020). Table S1 lists Z-scores and other information associated with all annotated genes mapped to strain BG2, and Table S2 lists Z-scores for data mapped to strain CBS138.

### Gene expression measurements by qRT-PCR

Replicate cultures of CBS138 and 2001 were grown to early log-phase in SCD medium at 30°C with shaking and then diluted to OD600 = 0.1. The cultures were divided and supplemented with micafungin (0.12 µg/mL) or vehicle control. After 1 hr of shaking, cells were pelleted and flash-frozen in liquid nitrogen. RNA was extracted using the hot acid phenol method (Pavesic et al. 2023), treated with DNAse I, and then cDNA was synthesized (Applied Biosystems cDNA Synthesis Kit cat. #4368814). Quantitative real-time PCR was performed on each sample using primers specific for *CDR1* or *FKS2* with ABsolute Blue qPCR Mix, SYBR Green (Cat. #AB4166) and a C1000 Touch thermal cycler (BioRad). The data were referenced to *PGK1* as an internal control in each sample.

### Growth assays

The *ira1Δ::URA3* mutants were grown separately to saturation in SCD medium, mixed at 1:1 cell ratio with their corresponding parent strains (BG2u, CBS138u, 2001u) that had been grown similarly, and then diluted 1000-fold into fresh medium. After 0 and 24 hr of regrowth, the mixed cultures were serially diluted, plated onto SCD agar medium and individual colonies were replica-plated onto SCD-ura medium to quantify percentage of mutant cells in the population. To ascertain fitness defects at elevated temperature, wild-type and mutant strains were cultured at 30°C to stationary phase in SCD medium, serially diluted in fresh medium (1:100 initial dilution followed by 1:5 dilutions), frogged onto SCD agar medium, and then incubated for 24 hr at 30°C or 37°C before photography. To quantify micafungin resistance, strains were grown to saturation in SCD medium, diluted into fresh medium containing varying doses of micafungin in the presence or absence of oligomycin (40 µg/mL), mixed, cultivated at 30°C without shaking, and then cell density was quantified by measuring OD600 using an Accuris Instruments SmartReader 96-T. IC50 was determined by fitting a sigmoid equation to the data and determining the drug dose at which half-maximal growth occurs. For checkerboard assays involving varying doses of two different compounds, the IC50s were determined for each compound separately and then the IC50 of the diagonal where both compounds are mixed at ∼1:1 ratio (in IC50 units) was determined. SDZ 90-215 was a kind gift from Novartis Pharmaceuticals. Myriocin was obtained from Cayman Chemicals. Manogepix was from Selleckchem.

## Data availability statement

All FastA files, site count files, gene count files, and genome feature files are available from the authors on request. All *C. glabrata* strains, plasmids, and pools of insertion mutants are available upon request. The authors affirm that all data necessary for confirming the conclusions of the article are present within the article, figures, and tables.

## ACKNOWLEDGMENTS

The authors thank Drs. Taiga Miyazaki, David Perlin, Erika Shor, Brendan Cormack, and Karl Kuchler for generously providing *C. glabrata* strains and helpful advice. We are grateful to Drs. John Kim and Andrew Gordus for providing access to critical instruments. Dr. Benoit Kornmann kindly suggested the “top-site removed” method for identification of passenger effects. Dr. Dominic Hoepfner (Novartis Biomedical Research) generously provided SDZ 90-215. Josh Schultz commented thoughtfully on the project and the manuscript. This research was supported by grants from the National Institutes of Health (T32-GM007231 to the JHU CMDB training program; R01-AI153414 to KWC).

